# USP48/USP31 Is a Nuclear Deubiquitinase that Potently Regulates Synapse Remodeling

**DOI:** 10.1101/2023.09.19.558317

**Authors:** Qi Ma, Hongyu Ruan, Huihui Dai, Wei-Dong Yao

**Author notes:** **Corresponidng author:** Wei-Dong Yao, Ph.D. SUNY Upstate Medical University E. Adams St Syracuse, NY 13210.

## Abstract

Deubiquitinases present locally at synapses regulate synaptic development, function, and plasticity. It remains largely unknown, however, whether deubiquitinases localized outside of the synapse control synapse remodeling. Here we identify ubiquitin specific protease 48 (USP48; formerly USP31) as a nuclear deubiquitinase mediating robust synapse removal. USP48 is expressed primarily during the first postnatal week in the rodent brain and is virtually restricted to nuclei, mediated by a conserved, 13-amino acid nuclear localization signal. When exogenously expressed, USP48, in a deubiquitinase and nuclear localization-dependent manner, induces striking filopodia elaboration, marked spine loss, and significantly reduced synaptic protein clustering in vitro, and erases ∼70% of functional synapses in vivo. USP48 interacts with the transcription factor NF-κB, deubiquitinates NF-κB subunit p65 and promotes its stability and activation, and up-regulates NF-κB target genes known to inhibit synaptogenesis. Depleting NF-κB prevents USP48-dependent spine pruning. These findings identify a novel nucleus-enriched deubiquitinase that plays critical roles in synapse remodeling.

## Introduction

Ubiquitination is a fundamental post-translational protein modification carried out by sequential actions of ubiquitin-activating enzymes (E1), ubiquitin-conjugating enzymes (E2), and substrate-selective ubiquitin ligases (E3), and plays crucial roles in nearly all eukaryotic processes (Hershko and Ciechanover, 1998). Ubiquitination is reversed by deubiquitinating enzymes (DUBs), a large family of proteases that cleave the isopeptide linkage between ubiquitin moieties and serve as a mechanism to control ubiquitination levels and ubiquitination-related functions (Komander et al., 2009; Mevissen and Komander, 2017). DUBs are divided into five classes of cysteine proteases, including ubiquitin-specific proteases (USPs), ubiquitin C-terminal hydrolases (UCHs), ovarian tumor proteases (OTUs), the Josephin family, and the motif interacting with ubiquitin-containing novel DUB family (MINDYs), and a family of the JAB1/MPN/Mov34 (JAMM) metalloproteases (Mevissen and Komander, 2017). USP DUBs, each containing a USP catalytic domain, are the largest cysteine subfamily protease with more than 50 members in humans.

Ubiquitination has emerged as a major mechanism in synapse development, function, and plasticity (Bingol and Sheng, 2011; DiAntonio and Hicke, 2004; Mabb and Ehlers, 2010; Tai and Schuman, 2008). Accumulating studies have revealed important roles for a number of DUBs in these processes (Goo et al., 2015; Kowalski and Juo, 2012; Widagdo et al., 2017). Specifically, ubiquitin C-terminal hydrolase (Ap-uch) is rapidly activated as an immediate-early gene required for long-term facilitation in *Aplysia* (Hegde et al., 1997). In mammals, UCH-L1 helps maintain synapse structure in cultured neurons in an activity-dependent manner, and is required for normal synaptic and cognitive function and, importantly, rescues synaptic and memory deficits in an Alzheimer’s disease mouse model (Cartier et al., 2009; Gong et al., 2006; Wood et al., 2005). Several DUBs in the USP family including USP6 (Zeng et al., 2019), USP8 (Bland et al., 2019; Kerrisk Campbell and Sheng, 2018; Scudder et al., 2014), USP9X (Yoon et al., 2020), USP14 (ATXN) (Wilson et al., 2002), USP46 (Huo et al., 2015; Kowalski et al., 2011) and CYLD (Colombo et al., 2021; Ma et al., 2017; Zajicek et al., 2022) have been reported to regulate various aspects of synapse formation and plasticity, glutamate receptor degradation and trafficking, and memory. Finally, the OTU family DUB A20 has recently been shown to negatively regulate synapse formation in hippocampal neurons in a NF-κB activation-dependent manner (Mei et al., 2020).

SynUSP (Synaptic USP), the rat ortholog of USP48 (formerly USP31) was first cloned from rat brain postsynaptic density and lipid raft fractions, suggesting a role in mammalian CNS neurons and synapses (Tian et al., 2003), although no evidence has since appeared to support this suggestion. In contrast, a number of studies have begun to reveal important functions of USP48 in NF-κB signaling (Li et al., 2018b; Liu et al., 2018; Schweitzer and Naumann, 2015; Tzimas et al., 2006), chromatin remodeling (Ji et al., 2015; Uckelmann et al., 2018), DNA damage response (Cetkovska et al., 2017), dopamine receptor control of blood pressure (Armando et al., 2014), and tumorigenesis in Cushing’s disease, glioblastoma and acute promyelocytic leukemia (Chen et al., 2018; Li et al., 2018a; Sbiera et al., 2019; Zhou et al., 2017). Of note, USP48 has been shown to interact with and stabilize the NF-κB subunit p65 and to promote NF-κB signaling (Liu et al., 2018; Schweitzer and Naumann, 2015). In neurons, NF-κB is constitutively active and plays essential and context-dependent roles in synapse formation during development and under pathologic conditions, as well as in synaptic plasticity and learning, largely via transcriptional modulation of various targeting genes (Kaltschmidt and Kaltschmidt, 2009; Meffert and Baltimore, 2005; Robison and Nestler, 2011).

In this study, we characterize the role of USP48 in synapse efficacy and remodeling. Contrary to the initial suggestion, we found that the full-length USP48 is predominantly localized in neuronal nuclei. We further show that nuclear USP48 potently inhibits synapse formation and limits synapse strength through the activation of NF-κB. Our work identifies a novel nuclear deubiquitinating mechanism that has profound effects in the remodeling of synapses.

## Results

### USP48 expression is virtually restricted to neuronal nuclei

We first characterized tissue and subcellular expression profiles of USP48 using immunoblotting (IB) and immunofluorescence (IF) assays (Figure 1). Compared to peripheral tissues, USP48 protein levels are markedly higher in several mouse brain regions including cortex, hippocampus, and cerebellum (Figure 1*A*). In the hippocampus, USP48 expression was the highest during the first postnatal week but maintained at a lower level thereafter (Figure 1*B, C*). This developmental profile is opposite to that of PSD-95, a postsynaptic scaffolding protein in excitatory neurons. Similar results were obtained in cultured neurons (Figure S1*A, B*). IB analysis of neuronal and glia cultures indicated that USP48 expression was more abundant in neurons (Figure 1*D*). Importantly, subcellular and nuclear fragmentation experiments demonstrated that USP48 expression was abundant in the nucleus, but undetectable in cytoplasmic and synaptic fractions (Figure 1*D, E*). Consistently, endogenous USP48 was primarily detected by IF in the nuclei of rodent hippocampal neurons in vivo and in vitro, although moderate expression was observed in cultured glia (Figure 1*G, H*, Figure S1*C*). Single-cell RT-PCR assay confirmed that USP48 and PSD-95 were expressed in the same neuron (Figure S1*D*). To further confirm the nuclear enrichment of USP48, we examined endogenous and GFP- or Myc-tagged USP48 in several mammalian cell lines. IF experiments confirmed that USP48 expression was virtually restricted to the nuclei of HEK293T, Neuro2a, and HeLa cells (Figure 1*I*; Figure S1*E-G*). Together, we conclude that USP48 is predominantly a nuclear DUB.

**Figure 1.**
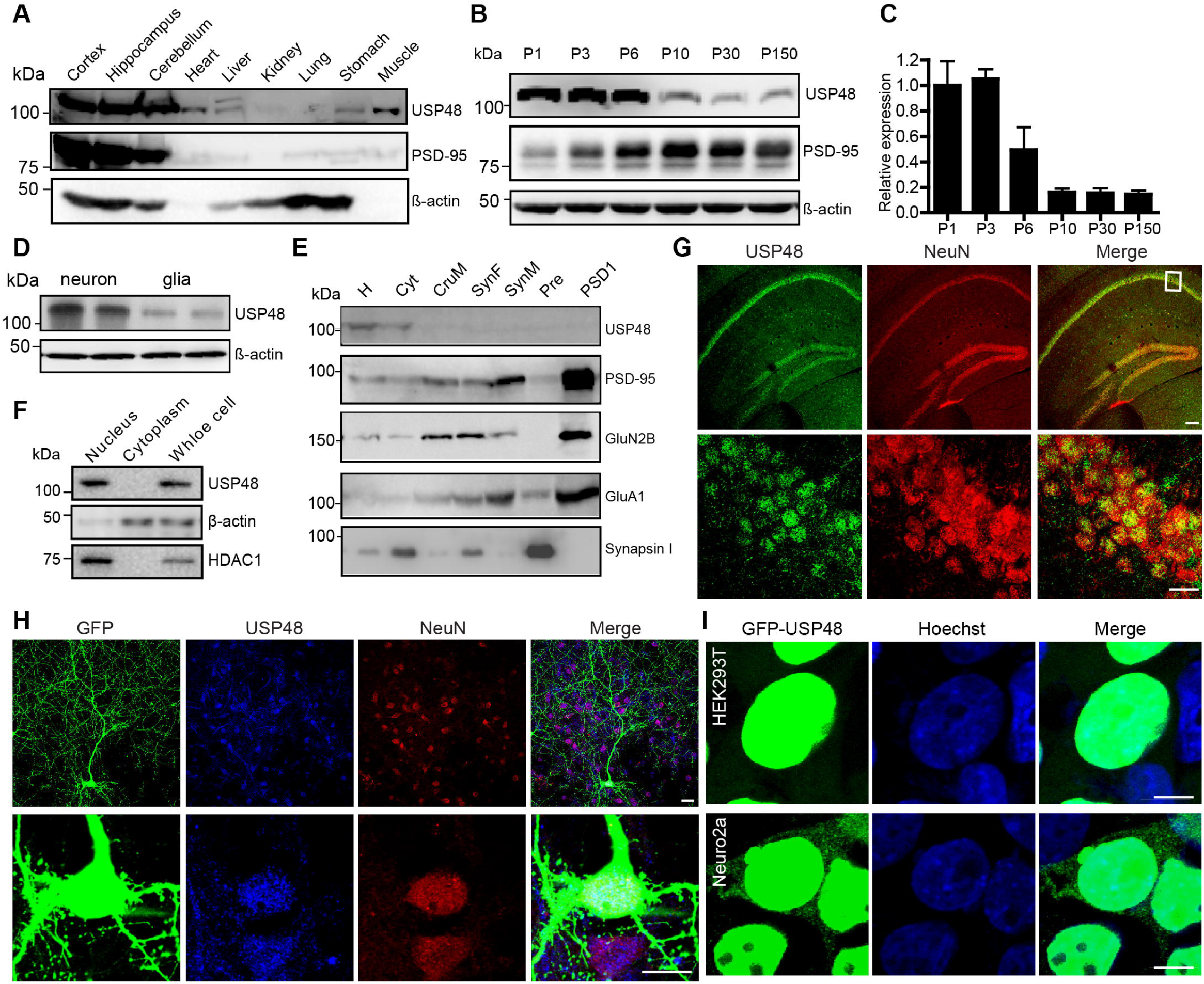
USP48 expression profiles and restricted localization to the nucleus. **A,** Immunoblotting (IB) analysis of USP48 expression in various mouse tissues. PSD-95 and β-actin are used as a neuronal marker and loading control, respectively. Note that β-actin was undetectable from heart and muscle, consistent with the fact that the most abundant actin in these tissues is α-actin. 20 µg total protein lysate from each tissue was loaded. **B, C,** Developmental expression of USP48 in the mouse brain. n = 3 independent experiments. **D,** Levels of USP48 expression in neurons and glia. **E,** IB analysis of subcellular fractions from the mouse brain. H, homogenates; Cyt, cytosol; CruM, crude synaptosomal membranes; SynF, synaptosome fraction; SynM, synaptic plasma membrane; Pre, synaptic vesicles enriched; PSD1, PSD fraction after one Triton X-100 extraction. **F,** IB of USP48, β-actin, and HDAC1 in nuclear and cytoplasmic fractions as well as total lysates prepared from mouse brains. **G**, IF images showing co-localization of USP48 (green) and the neuronal marker NeuN (red) on a three-month old mouse hippocampal section. Box (above) indicates where higher magnification images (below) were sampled. Scale bars, 50 µm (above), 20 µm (below). **H,** Co-localization of USP48 with NeuN in cultured rat hippocampus neurons. Scale bars, 20 µm. **I,** Nucleus-restricted GFP-USP48 in transfected HEK293T and Neuro2a cells. Scale bars, 10 µm.

### A nucleus localization signal in USP48

We next mapped the amino acid residues that are responsible for the nuclear localization of USP48. Full-length mouse USP48 consists of 1052 amino acids and possesses an N-terminal USP domain with conserved Cys- and His-boxes, three DUSP (domain in USPs) domains, and a ubiquitin-like (UBL) domain at the C-terminus (Figure 2*A*) (Tian et al., 2003). Cysteine 98 (C98) is responsible for the deubiquitination activity (Tian et al., 2003) and several phosphorylation sites (S903, S904, S905 and T940, together termed 3S1T) preceding UBL play regulatory roles (Schweitzer and Naumann, 2015). Myc-C98A and Myc-3S1T displayed similar nuclear localization as the full-length Myc-USP48. Myc-USP48 truncation mutants (Δ1, Δ3, Δ4, Δ5, and Δ7) lacking indicated amino acid stretches (Figure 2*A*) retained their nuclear preference but mutants lacking amino acids 796-946 (Δ2 and Δ6) lost the nuclear localization capability and instead were enriched in the cytoplasm (Figure 2*A, B*). Similar results were observed when Myc-USP48 and mutants were expressed in HEK293T cells (Figure S2*A*). We also obtained similar results using GFP-tagged USP48 and mutants (Figure S2*B*). These results suggested that amino acid residues within 796-946 were responsible for the nuclear localization of USP48.

**Figure 2.**
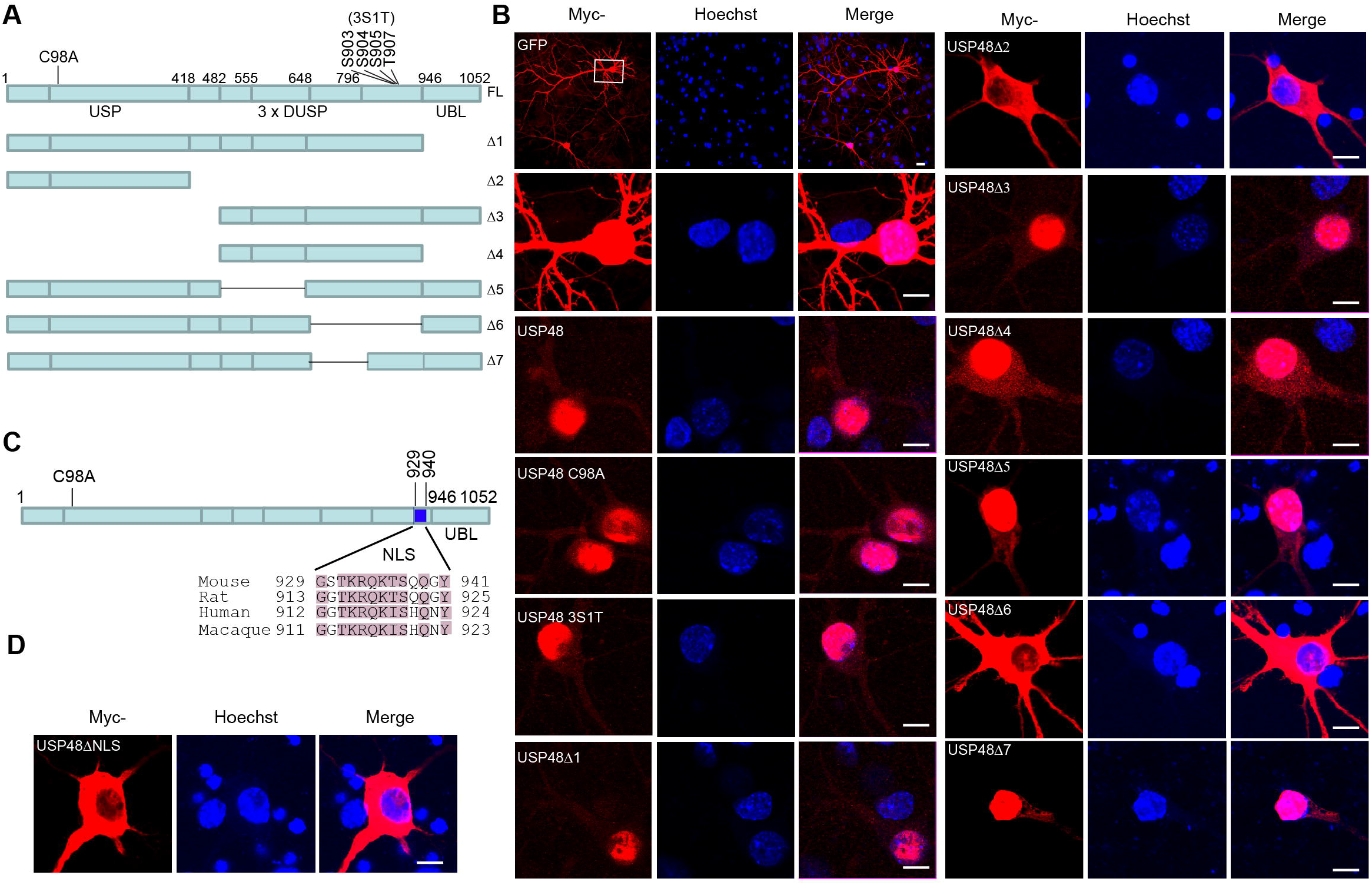
Mapping of nucleus localization motif in USP48. **A,** Schematic of full-length (FL) USP48 and mutants. Locations of C98 and phosphorylation sites (S903, S904, S905, T907) and USP, DUSP, and UBL domains are indicated. **B,** IF images of transfected cultured primary neurons showing subcellular localization of Myc-tagged GFP, wild-type full-length or mutant USP48 (red) relative to the nuclear Hoechst (blue) staining. Scale bars, 20 µm (above), 10 μm (below and all others). **C,** Alignment of the predicted NLS across different species. **D,** IF images of transfected neurons showing cytoplasm localization of Myc-USP48ΔNLS. Scale bar, 10 µm.

We reasoned that USP48 may contain a NLS and searched for this potential sequence using a NLS prediction software (cNLS Mapper) (Kosugi et al., 2009), which indeed predicted a putative 13-amino acid (929-941) motif (GSTKRQKTSQQGY) in the mouse USP48. This sequence is conserved among mice, rats, humans, and macaque (Figure 2*C*). Deleting this motif (Myc-USPΔNLS and GFP-USPΔNLS) virtually abolished the nuclear presence of USP48 and allowed the protein to localize primarily in the cytoplasm in cultured neurons (Figure 2*D*) and HEK293T cells (Figure S2*C*). These results identified a NLS motif that is necessary for localizing USP48 to the nucleus.

### Naturally existing shorter cytoplasmic variants of USP48

An NCBI database search for human and mouse USP48 transcripts and proteins revealed shorter isoforms lacking DUSP and UBL domains as well as the NLS motif in both species (Figure S3*A, B*). We thus obtained the human (hUSP48-S; available from Addgene) and cloned the mouse (mUSP48-S; Figure S3C) short USP48 isoforms from a mouse brain cDNA library and examined the subcellular localization of these naturally existing variants in cultured hippocampal neurons. As expected, both hUSP48-S and mUSP48-S displayed abundant localization throughout the cytoplasm including dendrite of neurons with minimal nuclear presence (Figure S3*D*). These data further support that the NLS motif identified above is necessary for the accumulation of the full-length USP48 to the nucleus.

### USP48 remodels dendritic spine morphologies and densities in cultured neurons

GFP-actin has been used to label neurons with enhanced structural details at dendrites and spines (Choidas et al., 1998; Fischer et al., 1998). To gain initial insights about the role of USP48 in synapse remodeling, we examined the effects of USP48 overexpression in GFP-actin labeled neurons. Cultured hippocampus neurons expressing GFP-actin alone displayed robust mushroom, stubby and thin spines with standard morphologies (Figure 3*A, B*). Strikingly, co-expression of USP48, but not the enzyme-dead mutant C98A, with GFP-actin produced a highly unusual phenotype characterized by a near loss or marked reduction of normal mushroom, stubby and thin spines, resulting in a dramatic decrease of total spines. This was accompanied with a significant increase of filopodia protrusions and the appearance of abundant atypically shaped filopodia (Figure 3*A-C*). To examine whether these phenotypes held true in the absence of GFP-actin, we repeated these experiments with DsRed (no actin) as a neuron fill. USP48 expression, but not C98A, still significantly reduced the total, mushroom and stubby spine densities, but did not induce the striking abnormal spine morphology (Figure 3*D, E*). Finally, expression of USP48 mutant lacking the NLS (Myc-USP48Δ6) failed to overly alter dendritic spine densities or morphologies (Figure 3*F-G*). Thus, nuclear, but not cytoplasmic, USP48 appears to control, in a deubiquitinase activity-dependent manner, dendritic spine numbers and shape, and likely through modulation of the actin cytoskeleton.

**Figure 3.**
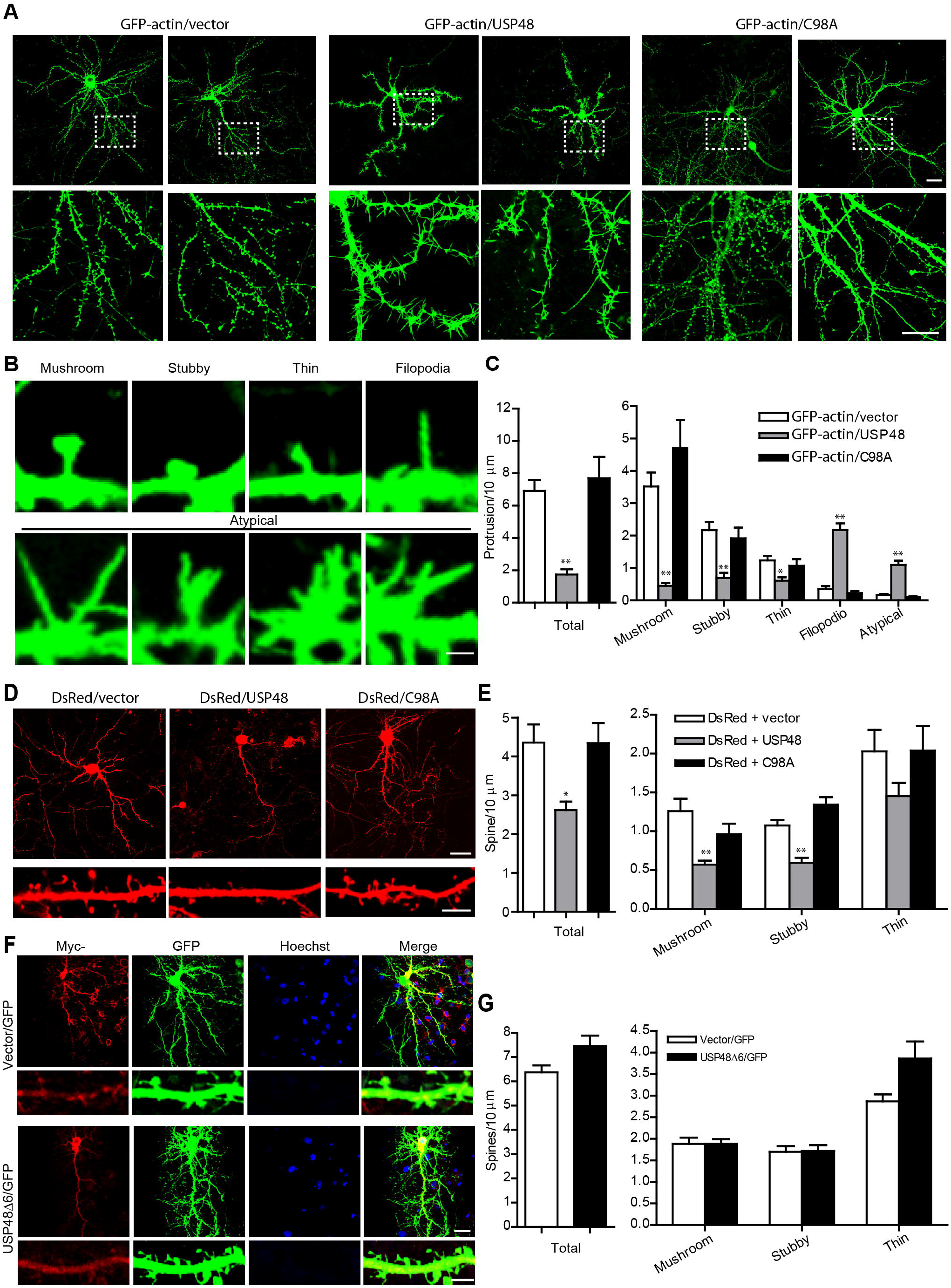
USP48 potently modulates dendritic spine morphogenesis. **A,** Representative neurons (above) and dendritic segments at higher magnifications (below) co-transfected with USP48, C98A, or vector control and GFP-actin. Boxes indicate where higher magnification images were derived. Scale bars, 40 µm (above), 25 µm (below). **B,** Mushroom, stubby and thin spines, filopodia and atypical structures observed from (**A**) at a higher magnification. Scale bar, 2 µm. **C**, Quantifications of (**A**) and (**B**). n = 16, 16, 15 dendrites in GFP-actin/vector, GFP-actin /USP48, and GFP-actin /C98A groups, respectively. *p < 0.05, **p < 0.01, one-way ANOVA with post hoc Dunnetts’ tests vs. vector control. **D,** Representative neurons and dendritic segments from neurons co-transfected with USP48 or C98A and DsRed. Scale bars, 20 µm (above), 5 µm (below). **E,** Quantifications of spine densities from **D**. n = 11, 10, 10 in DsRed/vector, DsRed/USP48, and DsRed/C98A groups, respectively. *p < 0.05, **p < 0.01, one-way ANOVA with post hoc Dunnetts’ tests vs. vector control. **F,** Representative fluorescence images of neurons and dendritic segments transfected with Myc-GFP/GFP or Myc-USP48Δ6/GFP. Hoechst labels nuclei (blue). Scale bars, 20 µm (above), 5 µm (below). **G**, Quantifications of (**F**). n = 18 and 14 for Myc-GFP/GFP and Myc-USP48Δ6/GFP groups, respectively.

We further investigated USP48 modulation of spine densities by gene knockdown and knockout (Figure 4). Using a shUSP48 hairpin (sequence “a”, targeting both full-length USP48 and USP48-S), we were able to silence the endogenous USP48 by ∼70% in HEK293T cells and ∼90% in cultured hippocampal neurons (Figure S4*A-D*). USP48 knockdown significantly increased total, mushroom, and stubby spine densities (Figure 4*A, B*). Importantly, the knockdown effects were prevented by co-expression of an RNAi-resistant USP48 (Figure 4*A, B;* Figure S4*E*). This result was further confirmed by a CRISPR/Cas9-mediated knockout strategy which completely deleted USP48 in stable HEK293T cells but removed about 70% of endogenous USP48 in cultured neurons at the time of our experiments (Figure S4*F-I*)). USP48 knockout significantly increased total as well as mushroom, stubby and thin spine densities (Figure 4*C, D*), which was reversed by co-expression of the RNAi-resistant USP48 (Figure 4*C, D*). Taken together, these results demonstrate that nucleus localized USP48 negatively regulates dendritic spine density and morphology.

**Figure 4.**
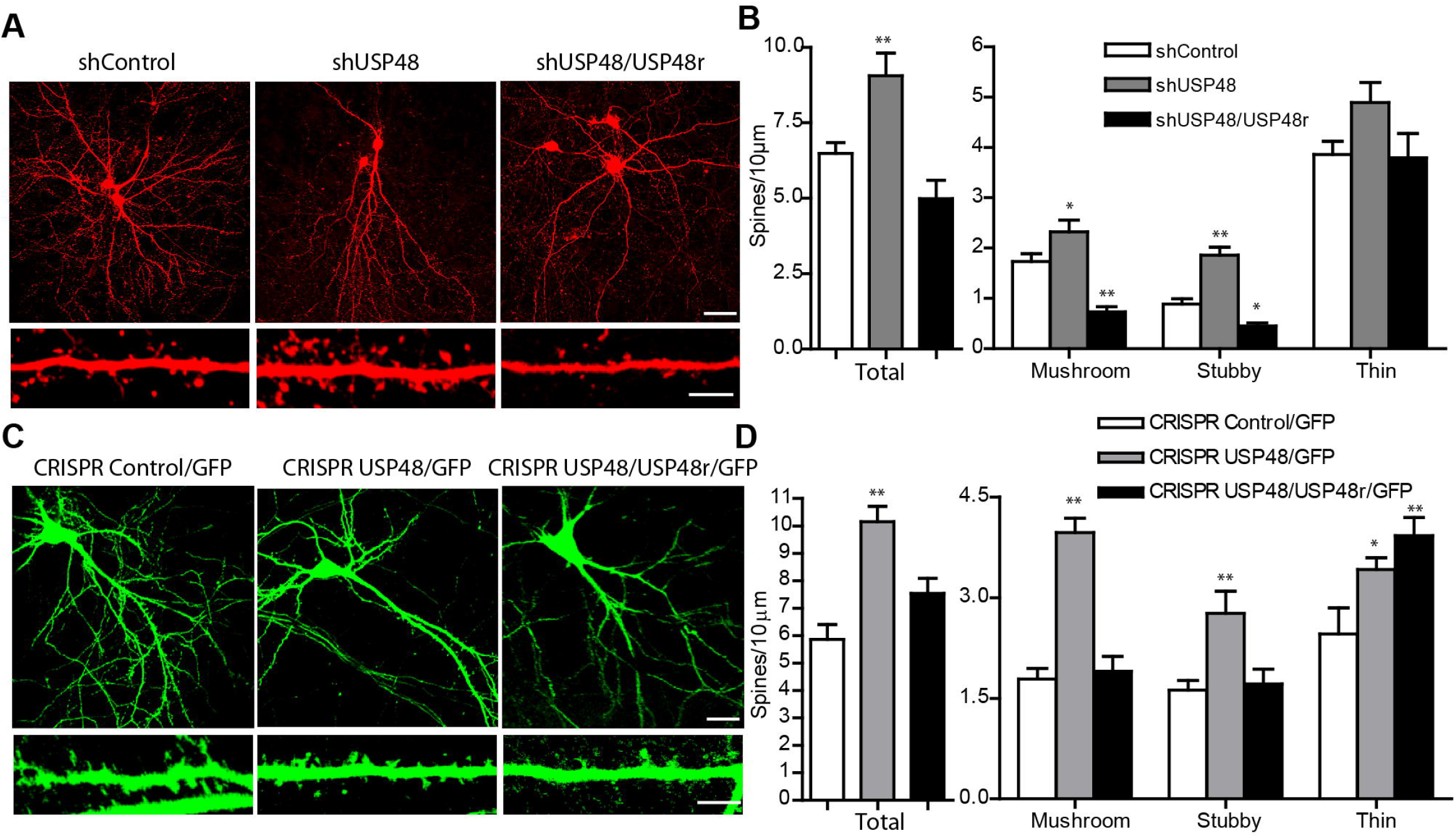
USP48 depletion promotes dendritic spine formation. **A**, Representative neurons and dendrites transfected with shControl, shUSP48, or shUSP48/USP48r. Scale bars, 20 µm (above), 5 µm (below). **B**, Quantifications of spine densities from (**A)**. n = 18, 18, 17 in shControl, shUSP48, or shUSP48/USP48r groups, respectively. *p < 0.05, **p < 0.01, one-way ANOVA with post-hoc Dunnett’s tests vs. shControl. **C,** Representative neurons and dendrites transfected or co-transfected with CRISPR/GFP, CRISPR USP48/GFP, or CRISPR USP48/USP48r/GFP. Scale bars, 20 µm (above), 5 µm (below). **D,** Quantifications of spine densities from (**C)**. n = 10, 13, 13 in CRISPR Control/GFP, CRISPR USP48/GFP, and CRISPR USP48/USP48r/GFP groups, respectively. *p < 0.05, **p < 0.01, one-way ANOVA with post hoc Dunnett’s tests vs. control.

### USP48 reduces synaptic protein levels and clustering

We next investigated the effect of USP48 on synaptic proteins. IB analysis indicated that total levels of glutamate receptors GluA1, GluA2, and GluN2A were significantly reduced, and GluN1, GluN2B and the presynaptic marker Synapsin I were also decreased (although the p-values did not reach a significance level) by overexpression of USP48 (Figure 5*A, B*). In comparison, knockdown of USP48 had largely opposite effects on levels of these proteins (Figure 5*C, D*). Biotinlyation experiments further showed that surface levels of GluA1 and GluA2 were also increased by USP48 knockdown (Figure S5*A, B*). Consistently, IF imaging indicated that USP48 overexpression and knockdown decreased and increased, respectively, the cluster intensity of surface GluA1 (sGluA1) (Figure 5*E, F, I*). Overexpression of full-length USP48, but not C98A or USP48-S, significantly reduced cluster intensity of Synapsin I (Figure 5*G, J*, Figure S5*C, D*). USP48, but not C98A overexpression also reduced the cluster intensity of PSD-95 (Figure S5*E, G*). In contrast, we found that knockdown of USP48 had largely opposite effects by increasing Synapsin I and PSD-95 clustering (Figure 5*H, J*; Figure S5*F, G*). Taken together, these results indicate that USP48 plays a significant role in the regulation of synapse strength.

**Figure 5.**
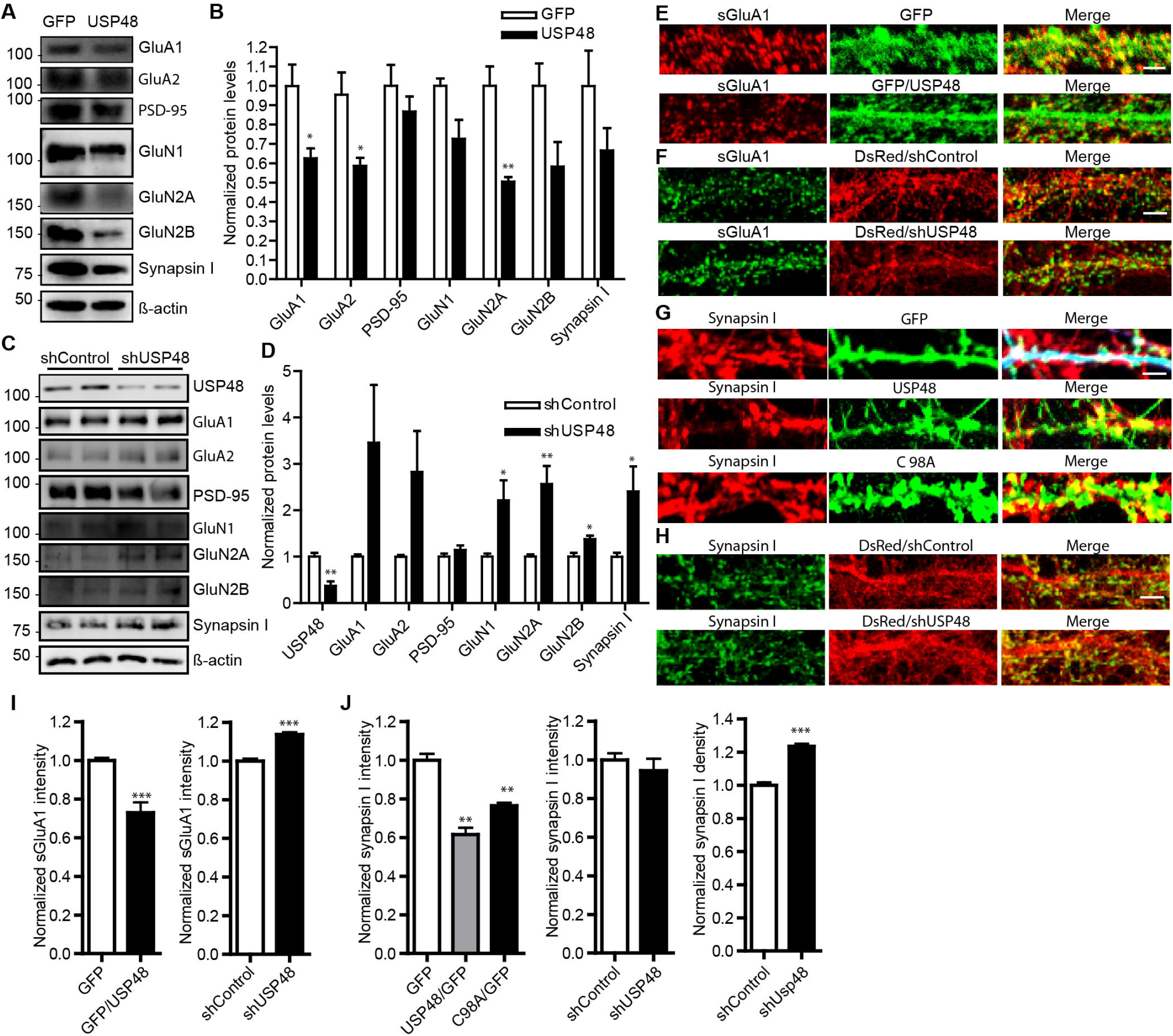
USP48 modulates levels and clustering of glutamatergic synaptic proteins. **A,** IB of indicated synaptic proteins in GFP- and USP48-overexpressing neurons. **B,** Quantifications of protein levels from (**A)**. n = 3 independent experiments. *p < 0.05, **p < 0.01; unpaired t tests. **C,** IB of indicated synaptic proteins in shControl- and shUSP48-expressing neurons. **D,** Quantification of protein levels from (**C)**. n = 4 independent experiments. *p < 0.05, **p < 0.01; unpaired t tests. **E, F,** Representative sGluA1 clusters from neurons infected with GFP or USP48/GFP (**E**), or with shControl or shUSP48 (**F**) lentivirus. Scale bars, 10 μm. **G, H,** Representative images of Synapsin I clusters from neurons transfected with GFP, USP48/GFP, or C98A/GFP (**G**), and neurons transfected with shControl or shUSP48 (**H**). Scale bars, 5 μm (**G**) and 10 μm (**H**). **I**, Quantifications of sGluA1 cluster intensity from (**E, F**). n = 16 and 16 dendritic branches (one branch/neuron) for GFP and USP48/GFP, and n = 53 and 54 dendritic branches for shControl and shUSP48, respectively. ***P < 0.001, unpaired t tests. **J**, Quantifications of Synapsin I cluster intensity and density from (**G, H)**. n = 22, 19 and 18 dendritic branches for GFP, USP48/GFP and C98A/GFP, respectively. **p < 0.01, one-way ANOVA with post hoc Dunnett’s test vs. control. Intensity: n = 24 and 21 in shControl and shUSP48 groups, respectively. Density: n = 32 for shControl and shUSP48 groups. ***p < 0.001; unpaired t tests.

### USP48 potently inhibits dendritic spine formation and function in vivo

To investigate whether USP48 remodels synapses in vivo, we injected viruses expressing GFP, USP48, C98A, shControl or shUSP48 into the mouse brain at different postnatal stages to manipulate the endogenous USP48 level, followed by dendritic spine imaging or slice electrophysiology (Figure 6). Compared to the GFP control, USP48, but not C98A lentivirus drastically reduced total dendritic spine density in both prefrontal cortex and hippocampus pyramid neurons (Figure 6*A, B, D*; Figure S6*A, B, D*). Consistently, shUSP48 lentivirus significantly increased dendritic spine density in these neurons (Figure 6*C, D*; Figure S6*A, C, D*). Similar results were also observed from AAV-mediated USP48 manipulations (Figure S6*E-H*). Importantly, USP48 overexpression in the PFC significantly decreased the frequency, but not amplitude, of mEPSCs recorded from infected pyramid neurons, indicating that that USP48 negatively regulates the number of functional synapses in vivo (Figure 6*E-G*). Taken together, these results suggest that USP48 strongly limits synaptic formation in vivo, likely independent of brain regions.

**Figure 6.**
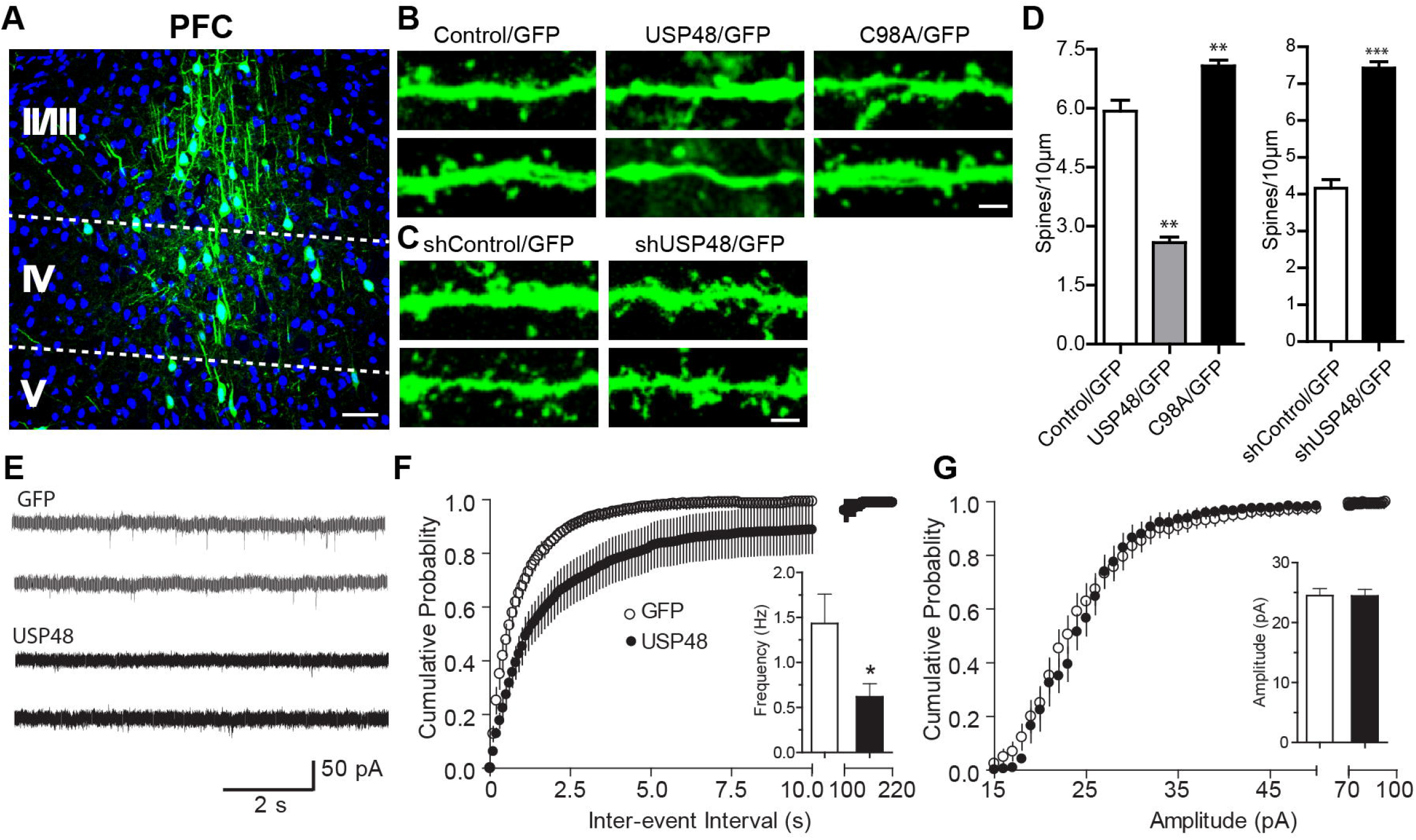
USP48 regulates dendritic spine density and synaptic strength in vivo. **A,** A representative coronal PFC section from the mouse prelmbic region showing lentivirus infected pyramidal neurons expressing USP48/GFP. Nuclei were labeled with Hoechst 33342 (blue). Cortical layers are labeled. Scale bar, 50 μm. **B**, Representative dendritic branches at higher magnifications showing individual spines from GFP, USP48/GFP, or C98A/GFP (**B**), and shControl/GFP or shUSP48/GFP (**C**) lentivirus infected neurons. Scale bars, 5 μm. **D**, Quantifications of total spine densities from (**B, C)**. n = 12, 12, 14 in GFP, USP48/GFP, and C98A/GFP groups, respectively. **p < 0.01, one-way ANOVA with post hoc Dunnett’s tests. N = 12 in shControl and shUSP48 groups. ***p < 0.001, unpaired t test. **E**, Representative mEPSC traces from AAV2-GFP or AAV2-USP48/GFP infected mouse PFC neurons. **F, G,** Cumulated probability plots of inter-event intervals (**F**) and amplitude (**G**) of mEPSCs from GFP and USP48/GFP neurons. Mean ± s.e.m. are shown as insets. n = 12 and 10 neurons for GFP and USP48/GFP, respectively. *p < 0.05, unpaired t test.

### USP48 deubiquitinates p65 and promotes its activation

Given its virtually restricted nuclear localization, we hypothesized that USP48 plays a role in important nuclear events. We focused on NF-κB signaling because of its involvement in synapse remodeling (Kaltschmidt and Kaltschmidt, 2009; Meffert and Baltimore, 2005) and its regulation by USP48 in non-neuronal systems (Liu et al., 2018; Schweitzer and Naumann, 2015). Co-immunoprecipitation analysis showed that USP48 interacted with p65 in the mouse brain and HEK293T cells (Figure 7*A*; Figure S7*A*). The deubiquitinating activity of USP48 was confirmed by biochemical reconstitution experiments using purified proteins in vitro (Figure S7*B,C*). In addition, we found that USP48, but not C98A cleaved K48-linked polyubiquitin chains on p65 in HEK293T cells (Figure 7*B*). Consistently, depletion of USP48 from cultured neurons (Figure 7*C*), Neuro2a cells (Figure S7*D*) and HEK293T stable cells (Figure S7*E*) increased K48 ubiquitination of p65. These results support previous studies that USP48 interacts with and deubiquitinates p65 in non-neuronal cells, and further demonstrate that this is also the case in neurons.

**Figure 7.**
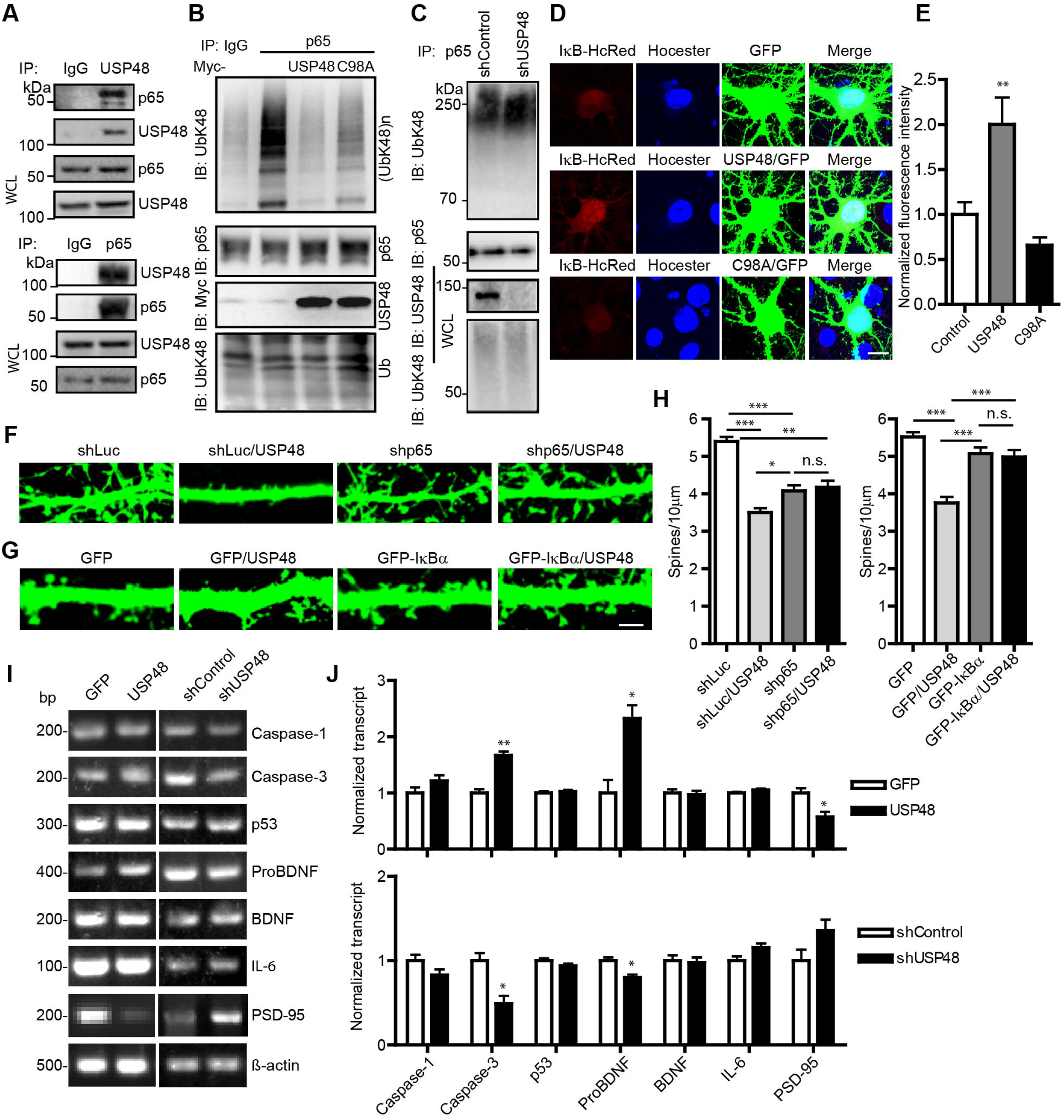
USP48 modulation of dendritic spine remodeling requires NF-κB signaling. **A,** Co-precipitation of USP48 and p65 from mouse brain lysates. IgG was used as a control. WCL, whole-cell lysates. **B,** Effects of Myc-USP48 or Myc-C98A in K48-linked polyubiquitination ofp65 in HEK293T cells. **C**, Effects of USP48 knockdown in K48-linked polyubiquitination of p65 in neurons. Ubiquitination of p65 in (**B**) and (**C**) was detected by anti-UbK48. **D,** Effects of GFP, USP48 or C98A overexpression on HcRed fluorescence intensity in a reporter gene assay for NF-κB activation in cultured neurons. Scale bar, 10 μm. **E,** Quantification of HcRed fluorescence from (**D)**. n = 9 neurons in each group. **p < 0.01; one-way ANOVA with post hoc Dunnett’s tests vs. control. **F,** Representative dendritic segments from shLuc, shp65, shLuc/USP48, or shp65/USP48 transfected neurons. Scale bar, 5 μm. **G,** Representative segments from GFP, GFP/USP48, GFP-IκB or GFP-IκB /USP48 transfected neurons. Scale bar, 5 μm. **H,** Quantification of total spine density from (**F)** and **(G)**. n = 15, 15, 16 and 16 neurons for shLuc, shp65, shp65/USP48 and shLuc/USP48 respectively. n = 14 neurons in each of the GFP control, GFP-IκB, GFP-IκB/ USP48 and GFP/USP48 groups. *p < 0.05, **p < 0.01, ***p < 0.001; one-way ANOVA with post hoc Turkey’s multiple comparison tests. **I,** Detections of indicated NF-κB target genes by RT-PCR after USP48 overexpression or knockdown in cultured neurons. β-actin was included as loading control. **J,** Quantification of relative gene expression from (**I)**. n = 3 independent experiments. *p < 0.05, **p < 0.01; unpaired t-tests.

Previous studies in non-neuron cells suggest that USP48 promotes NF-κB stability (Schweitzer and Naumann, 2015). Consistent with this idea, USP48, but not C98A enhanced the fluorescence intensity of nuclear IκB-HcRed, a fluorescence indicator of NF-κB activation (Mei et al., 2020; Wang et al., 2010), when co-expressed in cultured neurons (Figure 7D,E). Additional luciferase and fluorescence reporter assays further demonstrated that NF-κB activation was inhibited and enhanced, respectively, by USP48 depletion and overexpression in cultured neurons and HEK293T cells (Figure S7*F-H*). Taken together, these results support that USP48 stabilizes p65 and promotes its activation in neurons.

### USP48 remodeling of dendritic spines requires NF-κB

We next tested whether NF-κB mediated USP48-dependent dendritic spine remodeling. Knocking down p65 (by ∼80%, Figure S7*I, J*) resulted in a significant decrease of total spine density in cultured hippocampal neurons (Figure 7*F, H*), consistent with the requirement of p65 in synaptogenesis (Boersma et al., 2011). USP48 overexpression induced marked loss of spines, which was significantly more pronounced than that caused by p65 knockdown, and importantly, this effect of USP48 was prevented by knockdown of p65 (Figure 7*F, H*). Furthermore, inhibition of the endogenous p65 by introducing the exogenous GFP-IκBα (Boehm et al., 2007; Mei et al., 2020), an NF-κB inhibitor that binds p50 ⁄p65 dimer and prevents them from translocating to the nucleus, did not affect spine density by itself but prevented USP48-induced spine loss (Figure 7*G, H*). These results suggest that USP48 remodeling of dendritic spines is partially mediated by NF-κB.

Since USP48 promoted NF-κB activation, we further examined whether USP48 regulated expression of NF-κB target genes. Using reverse transcription-PCR, we examined transcript levels of a set of NF-κB target genes, including Caspase-1, Caspase-3, p53, ProBDNF, BDNF, IL-6, and PSD-95. We found that Caspase-3 and pro-BDNF were significantly upregulated while PSD-95 downregulated by USP48 overexpression, but other genes were not significantly altered (Figure 7*I, J*). In contrast, USP48 knockdown significantly reduced Caspase 3 and ProBDNF expression, without altering other genes (Figure 7*I, J*). These results indicate that USP48 selectively regulates expression of certain NF-κB target genes.

## Discussion

Here we identify a deubiquitinase that is primarily located in the nucleus of neurons but exerts potent constraint on dendritic spine formation and morphogenesis and synaptic efficacy both in vitro and in vivo. The nuclear localization of full-length USP48 is mediated by a conserved NLS, which is absent from a naturally existing, less abundant short isoform localized in the cytoplasm. We show that the profound spine remodeling effects of USP48 depend on its nuclear localization and deubiquitinase enzyme activity. Mechanistically, USP48 interacts with and deubiquitinates NF-κB, promoting its levels and activation in the nucleus and expression of certain target genes, which may, at least partially, mediates the effects of USP48 in synaptic remodeling. Our results identify the first DUB that is predominantly localized in neuronal nuclei but drives a potent nuclear program to globally regulate remodeling of distal synapses.

The highly abundant expression of full-length USP48 in brains and neurons, especially at early stages, suggests that it plays important roles in neural development and plasticity. Indeed, bi-directional manipulations of USP48 expression at early neuron development results in striking alterations in dendritic spine morphogenesis and synapse strength, suggesting that optimal levels of USP48 are crucial for proper maintenance of early synapse development and remodeling. Peak USP48 expression occurs during first postnatal week, which is before visible dendritic spine formation in the period leading to rapid synaptogenesis, consistent with a role of USP48 in limiting synapse formation. In comparison, the developmental profile of USP48 inversely correlates with that of PSD-95, which promotes synapse formation and maturation (El-Husseini et al., 2000) during peak synaptogenesis. Thus, a down-regulation of USP48 level may permit or facilitate the synaptogenesis process. Interestingly, USP48 has been shown to restrict cellular plasticity of differentiated cells in C. elegans (Rahe and Hobert, 2019). Finally, USP48 is one of the top upregulated genes in V1 layer 4 pyramidal neurons following visual deprivation-induced homeostatic scaling during critical period, though the functional significance of this is unknown (Steinmetz et al., 2016).

Our study identifies a unique DUB that is predominantly localized in neuronal nuclei yet possesses a role in remodeling distal synapses. The 13-amino acid NLS we have identified is necessary for retaining USP48 to the nucleus. This NLS is adjacent to a recently identified sequence in the rat USP48 (Liu et al., 2018), and it is possible that these sequences act together to regulate nucleocytoplasmic transport of USP48. Interestingly, a shorter, likely alternative splicing variant that lacks the carboxyl terminal NLS, exists naturally in both the mouse and human genomes. These “cytoplasmic” isoforms are likely much less abundant compared to the full-length isoform because our IF and IB analyses of endogenous USP48 rarely detect appreciable levels of cytoplasmic fluorescence signals. Importantly, the nucleus-enriched full-length USP48, but not USP48 mutant lacking the NLS, possesses the unique ability to remodel dendritic spines, suggesting that localization of USP48 to the nucleus and likely nuclear programs such as gene transcriptions are necessary to drive USP48-dependent synapse remodeling. Finally, it is worth noting that while the full-length USP48 is preferentially nucleus-localized, it is expected to present in the cytoplasm as well, albeit at very low abundance. Consistently, transcript for SynUSP, the rat ortholog of USP48, appears to be present in cortical PSD and lipid rafts fractions (Tian et al., 2003).

Recent studies in non-neuronal mammalian cell lines show that USP48 interacts with p65 (Liu et al., 2018; Schweitzer and Naumann, 2015), trims K48 polyubiquitin chains and confers stability to p65, thus presumably enhancing p65 signaling (Schweitzer and Naumann, 2015) (but see (Tzimas et al., 2006)). In this study, we have substantially extended those findings to neurons and shown that USP48 associates with p65 in the brain, negatively regulates K48 ubiquitination of p65 in cultured neurons, and potentiates p65 activation in neuronal nuclei. Neuronal NF-κB is localized at synapses and p65 translocates to the nucleus driven by constitutive basal synaptic activity or stimuli (e.g. depolarization and glutamate) mediated by synaptic Ca2+ signaling, thus transmitting synaptic signals to nuclear gene expression (Meffert et al., 2003). Nuclear p65 undergoes ubiquitination and subsequent proteasomal degradation, often in a DNA-bound form at certain target promoters (Bosisio et al., 2006; Colleran et al., 2013), which represents a major mechanism by which NF-κB signaling is terminated and the induced gene transcription is turned off. In non-neuronal cells, promoter bound-p65 ubiquitination has been reported to be antagonized by USP7/HAUSP, a nuclear DUB that is recruited to NF-κB target promoters following Toll-like- and TNF-receptor stimulation to prolong transcription of some inflammatory gene (Colleran et al., 2013). The full-length USP48 represents just the second nuclear DUB (among ∼100 putative DUBs in the human genome (Mevissen and Komander, 2017; Nijman et al., 2005)) that regulates nuclear NF-κB signaling. USP48 may regulate NF-κB target genes in a similar manner by deubiquitinates p65 bound to promoters of specific target gene sites. Our findings that a subset, but not all, of NF-κB target genes are bidirectionally regulated by USP48 overexpression or knockdown supports this possibility. Finally, neuronal NF-κB displays the highest basal activity before synaptogenesis, which coincidently parallels that of USP48 expression, and it has been suggested that posttranslational modification may be responsible for its activation (Boersma et al., 2011; Kaltschmidt and Kaltschmidt, 2009). It will be interesting to investigate whether USP48-dependent deubiquitination contributes to constitutive NF-κB activity in neurons and fine-tunes NF-κB dependent gene expression during early neuronal and synaptic development.

Our study reveals a DUB-mediated nuclear mechanism that bidirectionally drives synapse remodeling: increasing the full-length USP48 expression reduces spine density, inhibits spine maturation, and markedly decreases functional synapse numbers and weakens synaptic efficacy in vivo, whereas decreasing USP48 expression has largely opposite effects. Importantly, these synapse remodeling effects of USP48 depends on its deubiquitinase activity and nuclear localization, and are partially mediated by nuclear NF-κB. NF-κB can either positively or negatively regulate dendritic plasticity and synaptogenesis depending on developmental stages, cellular context, and subcellular localization (Boersma et al., 2011; Dresselhaus et al., 2018; Kishi et al., 2016), likely due to context-dependent regulation of different sets of target genes. Our observations that endogenous p65 level is required for normal synapse formation but USP48 overexpression strikingly inhibits synapse formation in a p65-dependent manner can be explained by USP48-dependent up-regulation of a subset of NF-κB target genes, including pro-BDNF and Caspase-3. Both genes have been reported to promote dendrite and synapse pruning (D’Amelio et al., 2012; Erturk et al., 2014; Guo et al., 2016; Koshimizu et al., 2009). USP48 downregulation of PSD-95 is somewhat surprising because NF-κB has been shown to turn on PSD-95 transcription (Boersma et al., 2011). This apparent inconsistency may suggest that other nuclear mechanisms may be engaged by USP48, in addition to NF-κB, to regulate PSD-95 gene expression. Finally, it is worth noting that USP48 appears to have its most striking effects in synapse remodeling in the presence of the exogenously expressed GFP-actin, which itself does not have overt effects in spine density and morphologies and in fact is routinely used as a dendritic spine label. The underlying mechanism is unknown but likely a result of USP48-dependent regulation of spine actin cytoskeleton which is unmasked in the presence of excessive actin.

Emerging studies have begun to reveal essential roles for USP48 in the pathogenesis of an increasing number of human diseases. By regulating NF-κB activation, genome stability, and intracellular signaling, mutations of USP48 have been reported to cause Cushing’s disease (Chen et al., 2018), glioblastoma (Zhou et al., 2017), and Fanconi anemia (Zhou et al., 2017). The human USP48 gene is mapped to a PARK6 locus (encoding the gene PINK1) associated with autosomal recessive Parkinson’s disease (Lockhart et al., 2004). Our findings that USP48 is a major nuclear DUB and potently regulates synapse remodeling and efficacy raise the possibility that this unique DUB may play a role in neural circuit assembly, plasticity and pathologies associated with some neurodevelopmental and neurologic diseases. In addition, USP48 associates with histone-marked chromatins, especially H3K27ac and H3K4me3 in mouse embryonic stem cells, and has been proposed as a histone modifier and chromatin eraser in epigenetic reprogramming (Ji et al., 2015). Thus, USP48 likely has multiple functions by targeting diverse substrates to regulate gene expression and remodel synapses, neurons, and circuits during development, plasticity, and maintenance.

## Materials and Methods

### Animals

Wild-type C57BL/6J mice were bred in house under standard conditions. Timed pregnant rats were purchased from Charles River Laboratories. Animal care and use protocols were approved by the Institutional Animal Care and Use Committee (IACUC) of State University of New York, Upstate Medical University.

### Antibodies

The following commercial antibodies were used: anti-Flag (Sigma, F7425), anti-Flag (Sigma, F3165), anti-HA (12CA5) (Roche, 11583816001), anti-c-Myc (9E10) (Santa Cruz, sc-40), anti-GST (Santa Cruz, sc-138), anti-GFP (Santa Cruz, sc-8334), anti-PSD-95 (Cell Signaling, 2507), anti-PSD-95 (Santa Cruz, #32290), anti-Synapsin I (Millipore, #1543), anti-GluN1 (Millipore, 05-432), anti-GluN2A (Millipore, 07-632), anti-GluN2B (Millipore, 06-600), anti-GluA1-NT (Millipore, MAB2263), anti-GluA2 (Millipore, MAB397), anti-NeuN (Millipore, MAB377), anti-GFAP (Cell Signaling, #3670), anti-USP48 (Abcam, ab72226), anti-USP48 (Abcam, ab62529), anti-HDAC1 (Cell Signaling, 5356), anti-ß-actin (Santa Cruz, sc-47778), anti-NF-κB p65 (Cell Signaling, #8242), anti-Ubiquitin (Santa Cruz, sc-8017), anti-Ubiquitin, Lys48-Specific (Millipore, #05-1307), Alexa Fluor goat anti-mouse IgG 488 (Invitrogen, A1100), 568 (Invitrogen, A-11004), 647 (Invitrogen, A21236), goat anti-rabbit IgG 488 (Invitrogen, A-11008) and 568 (Invitrogen, A-11036), and Horseradish peroxidase-conjugated goat anti-mouse (Pierce, #31430) and anti-rabbit (Pierce, #31460).

### Plasmids

pEGFP-USP48 was generated by sub-cloning the mouse USP48 into the mammalian expression vector pEGFP-C2 (Clontech). pCMV-Myc-USP48, pCMV-Myc-USP48 C98A, pCMV-Myc-USP48 S903A/S904A/S905A/T907R (pCMV-Myc-USP48 3S1T), pCMV-Myc-USP48Δ1, pCMV-Myc-USP48Δ2, pCMV-Myc-USP48Δ3, pCMV-Myc-USP48Δ4, pCMV-Myc-USP48Δ5, pCMV-Myc-USP48Δ6, pCMV-Myc-USP48Δ7 and pCMV-Myc-USP48ΔNLS were generated by sub-cloning the mouse USP48 and specific mutants into the pCMV-Myc vector (made in authors’ lab). Lentivirus transfer plasmids pLL3.7 EGFP-Myc-USP48, pLL3.7 EGFP-Myc-USP48 C98A, pLL3.7 EGFP-Myc-USP48 S903A/S904A/S905A/T907R (pLL3.7 EGFP-Myc-USP48 3S1T), pLL3.7 EGFP-Myc-USP48Δ1, pLL3.7 EGFP-Myc-USP48Δ2, pLL3.7 EGFP-Myc-USP48Δ3, pLL3.7 EGFP-Myc-USP48Δ4, pLL3.7 EGFP-Myc-USP48Δ5, pLL3.7 EGFP-Myc-USP48Δ6, pLL3.7 EGFP-Myc-USP48Δ7 and pLL3.7 EGFP-Myc-USP48ΔNLS were generated by sub-cloning the mouse USP48 and specific mutants into the pLL3.7 vector. The above USP48 and mutants were also sub-cloned into Syn-GFP-Syn-Myc-GFP or Syn-DsRed-Syn-Myc-DsRed vectors (made in authors’ lab) to drive expression in neurons. IκBα-HcRed was also sub-cloned into the Syn-GFP-Syn-Myc-GFP vector. Recombinant adeno-associated virus transfer plasmids pAAV-Syn-Myc-USP48 was generated by sub-cloning mouse USP48 and mutants into the pAAV-Syn-Myc-GFP vector. pLL3.7 GST-Myc-GFP and pLL3.7 GST-Myc-USP48 were generated by sub-cloning GST-Myc-GFP or GST-Myc-mouse USP48 into the pLL3.7 vector. DUAL hybrid cDNA Library -mouse cortical neurons (Bulldog Bio, P12401) and reverse primer: 5’ tcccagaatatcttacagacc 3’ were used to clone the mouse USP48 short isoform by PCR. CRISPR USP48 was constructed by cloning designed DNA sequences coding guide RNA (gRNA) of USP48 into LentiCRISPR (Shalem et al., 2014). Lok1-5 primer was used for sequencing and identification of CRISPR recombinants by PCR. USP48 gRNA 5’ ggagacggtgcggcccgagg 3’ targeting human, rat and mouse were designed based on the *Rattus norvegicus* Gene, Usp48 ENSRNOG00000013602, exon 1 (CDS: 45∼64bp). NF-кB-Luciferase and TK-Rellina have been described previously (Ma et al., 2008; Zhou et al., 2010). IκBα-HcRed was from Philip E. Auron. HA-tagged Ubiquitin (HA-Ub) and mutants were from Zhijian James Chen. pLLox3.7 was from Roger Nicoll. FUGW-DsRed or GFP was from Pavel Osten. Flag-HA-USP48 (# 22585) and GFP-p65 (# 23255) were from Addgene. All constructs were confirmed by DNA sequencing and protein expression.

### Cell culture

HEK293T, HeLa, Neuro2a, and primary glia cells were cultured in Dulbecco’s modified Eagle’s medium (DMEM) (Thermo Fisher Scientific, #11995073) supplemented with 10% fetal bovine serum (FBS) (GIBCO, #26140079). Cells were plated on 10 cm plates or 24 well plates with or without coverslips and grown to 50∼80% confluency before transfection. Hippocampal neurons were prepared from rat E18 embryos and maintained in Neurobasal Medium supplemented with B-27 and 2 mM L-glutamine (Invitrogen) for 18-21 days in vitro (DIV).

### Immunoprecipitation (IP) and immunoblotting (IB)

HEK293T cells were transfected with indicated plasmids with GenJet Transfection Reagent (SignaGen Laboratories), and at 48 hr cells were lysed in ice-cold RIPA buffer, consisting of 50 mM Tris-HCL pH 7.4, 0.1% SDS, 1% NP-40, 0.25% deoxycholic acid, 150 mM NaCl, 1 mM EDTA, 1 mM PMSF, and protease inhibitor cocktail (Roche). Neuro-2a cells, infected cultured neurons and mouse brain were similarly lysed/homogenized and cleared in ice-cold RIPA buffer. Samples were pre-cleared with protein A/G beads (Santa Cruz, sc-2003) at 4°C for 1 hr, followed by incubation with primary antibodies at 4°C for 2 hr. After incubation with protein A/G beads at 4°C overnight, immunocomplexes were washed with ice-cold RIPA buffer and eluted with SDS loading buffer by boiling for 5 min. Supernatant was collected by centrifugation and used in IB experiments. Proteins were resolved by SDS-polyacrylamide gel electrophoresis (SDS-PAGE) and transferred to a PVDF membrane (Bio-Rad). PVDF membranes were incubated with primary antibodies at 4°C overnight, followed by incubation with secondary antibodies at room temperature for 1 hr. Immunoreactivity signals were detected with the SuperSignal West Femto Maximum Sensitivity Substrate kit (Pierce) and a Chemico image system (Bio-Rad). Densitometry quantification of protein band intensities was carried out using ImageJ.

### Construction of stable cell lines and CRISPR/Cas9-mediated USP48 knockout in stable cells and neurons

HEK293T cells were transfected with LentiCRISPR or LentiCRISPR USP48 with Lipoamine 3000 (Invitrogen) following the manufacturer’s instruction. After 24 hr and then daily, cells were fed with fresh medium (DMEM plus 10% FBS and 400 µg/ml puromycin). Following screening for about one week, single clones would form and individual clones were transferred to new 24-well culture plates filled with fresh DMEM media plus 10% FBS (one clone/well) using pipette tip wetted with 0.25% trypsin. When the clones grew to 90% confluence, the cells were passaged for at least two generations. Stable cells were confirmed by immunoblotting, immunofluorescence and genome sequencing. Cultured neurons were co-transfected with Syn-GFP-Syn-Myc-GFP and indicated CRISPR constructs at DIV7.

### RNA interference

Mouse or rat USP48 shRNA target sequences: (a) 5’-gcatcttcagtacttgtttgc-3’, (b) 5’-gctctttgttgtggataaagt-3’, or (c) 5’-gctgatgcgtttcgtgtttga-3’ was cloned into FUGW-DsRed or -EGFP vector. Rat p65 shRNA target sequence: 5’-gagcatcatgaagaagagt-3’ was cloned into pLL3.7 vector. FUGW-EGFP and FUGW-DsRed containing irrelevant target sequences or shLuc was used as a negative control. These constructs were used to package lentivirus or rAAV along with helper vectors, or for transfection, where indicated.

### Virus preparation and infection

For lentivirus production, HEK293FT cells were co-transfected by calcium phosphate with an indicated transfer plasmid plus two helper vectors, PSHX2 and PMDG2. Supernatants were collected at 48 hr after transfection and centrifuged to concentrate the virus. To infect cultured neurons, concentrated virus solutions were added to DIV4∼7 hippocampal neurons and grown to DIV15∼21 before IB or IF and imaging analyses. For co-infection, two equal titers concentrated viruses were mixed thoroughly and added to DIV4 neurons. About 100% efficiency of co-infection was routinely obtained. For AAVs production, AAV-293 cells were co-transfected by polyethylenimine with an indicated transfer plasmid plus helper vector pDP2rs. AAVs were purified by serial centrifugations. AAV infection of neurons was done similarly as lentivirus.

### Biotinylation

Cultured hippocampal neurons at DIV20 (infected with indicated virus at DIV7) were live fed with EZ-Link Sulfo-NHS-SS-Biotin diluted in cold ACSF for 10 min on ice, washed three times with ice-cold ACSF, and lysated with ice-cold RIPA buffer. Lysates were cleared by centrifugation, and the suspension was incubated with NeutrAvidin Agarose at 4°C for 2 hr. After wash, proteins bound to NeutrAvidin Agarose were eluted into protein loading buffer for Western blotting assays. More procedural details are provided in the manufacturer’s manuals for EZ-Link™ Sulfo NHS-SS Biotinylation Kit (Pierce, 21445).

### Subcellular fractionation

Subcellular fractionation of mouse brains was performed as previously described (Ma et al., 2017). Fractions of whole cells, cytoplasm, and nucleus were obtained by centrifugation and resolved by SDS-PAGE and analyzed by IB.

### Ubiquitination and deubiquitination assays

HEK293T cells were collected and washed in ice-cold PBS 48 hr after transfection and lysed by denaturing 1% SDS RIPA buffer (containing 1 mM PMSF and protease inhibitor cocktail). Cell lysates were boiled at 100°C for 5 min and diluted with SDS-free RIPA buffer (final SDS concentration was 0.1%). Diluted lysates were homogenized and cleared by centrifugation (at >12000 rpm) at 4°C for 20 min. Total proteins in suspensions were quantified using the Bio-Rad Dc Protein Assay kit. Lysates were normalized against substrate proteins by IB before IP and IB analysis of target ubiquitination. Following IP with indicated antibodies, IB was done with indicated antibodies, and after stripping (with One Minute Western Blot Stripping Buffer, GM Biosciences, GM6001), re-probed with indicated antibodies for quantification of the amount of total proteins in the precipitates. Ubiquitination assays in Neuro-2a cells and infected cultured neurons were similarly done. For the reconstitution of deubiquitination reactions, HA-tagged Ub K48 was overexpressed in HEK293T cells and purified by IP with anti-HA under denatured conditions. GST-Myc-GFP or GST-Myc-USP48 was overexpressed in HEK293T cells and purified following modified instruction of Glutathione Sepharose 4B (GE Healthcare Life Sciences, 17-0756-01) using 1% NP40 buffer. Purified HA-Ub K48 was incubated with GST-Myc-GFP or GST-Myc-USP48 at 37°С for 4 hr following a deubiquitination procedure. IB was done with indicated antibodies to detect deubiquitination. More detailed procedures have been described previously (Ma et al., 2017).

### Luciferase assay

Luciferase-based reporter gene assays were performed using Dual Luciferase Reporter Assay System (Promega, E1910) as described previously (Ma et al., 2008; Mei et al., 2020).

### RT-PCR and single cell PCR

For RT-PCR, total RNA was extracted from infected hippocampus neurons by PureLink RNA Mini Kit following the manufacture’s instruction (Life technology). cDNA was synthesized using the extracted total RNA as a template and OligodT20 as a primer by SuperScript® III First-Strand Synthesis System (Invitrogen) following the manufacturer’s instruction. cDNA was used as the template for PCR amplification. For single cell PCR, individual neurons were patched with a patch electrode and contents of the cell were extracted into a PCR tube containing RT buffer and RNase inhibitors. cDNA was synthesized using Sensiscript RT Kit (QIAGEN) following the manufacturer’s instruction. cDNA was then used as the template for PCR amplification of specific genes.

### Neonatal intracerebroventricular (ICV) injections

AAVs expressing Myc-GFP, Myc-USP48 (and GFP), Myc-C98A (and GFP), shControl/GFP or shUSP48/GFP were injected bilaterally with a syringe into each lateral ventricle of cryoanesthetized P0 mice. At a desired age after injection, mice were used for electrophysiology recording or dendritic spine imaging.

### Stereotaxic virus injection, histology, and analysis

Lentivirus or AAVs expressing Myc-GFP, Myc-USP48 (and GFP), Myc-C98A (and GFP), shControl/GFP or shUSP48/GFP were stereotaxically injected bilaterally into the hippocampus of anesthetized P20 mice. The coordinates for hippocampal injections were (from Bregma): anteroposterior, -0.27; mediolateral, ±0.26; dorsoventral, -0.16 mm. The coordinates for PFC injections were: anteroposterior, +1.8 mm; mediolateral, 0.5 mm; dorsoventral, −2.5 mm. Two weeks after surgery, mice were euthanized and their brains were collected and fixed. Coronal sections (100∼150 μm) were cut for dendritic spine imaging and analysis. Sections were mounted on glass slides. For spine analysis, confocal images of Z stacks of ∼10 μm were acquired using a Leica SP5 confocal microscope fitted with a 40x oil objective at pixel resolutions of 512×512. Numbers of total spines (identified as GFP-filled protrusions) per 10 μm apical dendrites were measured from GFP-labeled pyramidal neurons infected with AAVs expressing Myc-USP48, Myc-C98A, shControl/GFP or shUSP48/GFP.

### Immunofluorescence (IF) and confocal microscopy

Cells were fixed in 4% paraformaldehyde (plus 5% sucrose for neurons) in PBS for 15 min, permeabilized with 0.3% Tween-20 in PBST for 20 min (skipped for morphology imaging and analysis), and blocked with 1% BSA for 30 min. Coronal brain sections (∼40 µm) were cut using a Microtome, permeabilized with 0.3% Triton X-100 in PBS, and blocked with 1% BSA for 1 hr. Cells or brain sections were incubated with indicated primary antibodies at 4°C for 3 hr or overnight, followed by incubation with goat anti-mouse or goat anti-rabbit secondary antibodies conjugated with appropriate Alexa dyes for 1 hr. For sGluA1labeling, cells were fed with anti-GluA1-NT antibody for 30 min in a 5% CO2 incubator, washed, fixed in 4% paraformaldehyde and 5% sucrose in PBS, and incubated with Alexa Fluor 568 second antibody. Cells or brain sections were mounted on glass slides for imaging.

For dendritic spine analysis, GFP- or DsRed-expressing dendrites (one or two segments per neuron) from DIV19-20 neurons transfected with indicated plasmids were traced using NeuronStudio software. Spine types including mushroom, stubby, and thin were automatically classified by NeuronStudio using the default setting and criteria implemented in the software. Protrusions that were not classified as spines were assigned manually as filopodia or atypical processes. Numbers of total, mushroom, stubby, and thin spines as well as filopodia and atypical protrusions per 10 μm were calculated. To quantify the intensity or density of sGluA1, Synapsin I, and PSD-95 clusters in cultured rat hippocampal neurons infected or transfected with indicated constructs, the fluorescence intensity or number of carefully traced clusters on randomly selected dendrites were measured. One or two dendrites were selected from individual neurons and the average fluorescence intensity or density of clusters was calculated for each segment before a group mean was derived.

For confocal microscopy, image stacks were acquired along the z-axis using a TCS SP5 Confocal Microscope (Leica Microsystems) fitted with a 40x objective (NA 1.4) at pixel resolutions of 512 x 512 or 1024 x 1024. Same imaging settings were applied for all scans when fluorescence intensity was compared. All groups to be compared were also run simultaneously using cells from the same culture preparations and transfection conditions. When measuring fluorescence intensity (pixel intensities), outlines of selected areas were traced and fluorescence signal was measured for the traced areas at desired channels. Background was subtracted before fluorescence intensity measurement as necessary. Images were analyzed using ImageJ. Means from multiple cells or dendritic segments in the same group were averaged to obtain a population average and presented as mean ± s.e.m. More detailed procedures have been described previously (Ma et al., 2017; Mei et al., 2020).

### Electrophysiology

Whole-cell patch-clamp recordings were performed on infected mouse pyramidal neurons at P20∼P23 after infection at P0 with AAVs expressing Myc-GFP or Myc– USP48/GFP. Pyramidal neurons were identified by their morphology. Cells were superfused at room temperature with an oxygenated (95% O2 and 5% CO2) extracellular solution containing (in mM) 126 NaCl, 25 NaHCO3, 11 glucose, 2.5 KCl, 1.2 NaH2PO4, 1.2 MgSO4, and 2.5 CaCl2. Recording pipettes (4.5–5.5 MΩ) were filled with an intracellular solution containing the following (in mM): 142 Cs-gluconate, 8 NaCl, 10 Hepes, 0.4 EGTA, 2.5 QX-314 [N-(2,6-dimethylphenylcarbamoylmethyl) triethylammonium bromide], 2 Mg-ATP, and 0.25 GTP-Tris (pH 7.25) (with CsOH). mEPSCs were recorded at −60 mV with picrotoxin (50 μM) and TTX (1 μM) present in the superfusion medium. All electrophysiological data were acquired using an Axon Multiclamp 700B amplifier (Molecular Devices) with Digidata 1440A and pClamp software (version 10; Molecular Devices). Signals were filtered at 1 kHz and digitized at 10–20 kHz. mEPSCs were analyzed by Mini Analysis 6 (Synaptosoft). All recordings were performed with the electrophysiologist blinded to the groups.

### Statistics

All data were presented as mean ± s.e.m. Unpaired Student’s t tests were used to compare two independent variables. One-way ANOVA with indicated post-hoc tests were used to compare multiple variables. Statistical significance was set at 0.05.

## Acknowledgements

We thank members of the laboratory for comments and discussions. This study was supported by NIH grants, MH106489, NS093097, DA032283, and RR026761 (W.-D.Y.).

## Author Contributions

Q.M., H.R. and W.-D. Y. designed research. Q.M, H.R and H.D. performed experiments and analyzed data. Q.M., H.R. and W.-D. Y. wrote the paper. The authors declare no conflict of interest.

