## Supplemental figure legends for "USP48/USP31 Is a Nuclear Deubiquitinase that Potently Regulates Synapse Remodeling"

**Supplementary Legends**

**Figure S1. Additional data on USP48 expression profile and nuclear enrichment. A,** IB analysis of USP48 protein levels at different days in vitro (DIVs) in cultured rat hippocampus neurons. **B**, Quantification of relative USP48 levels from (**A)**. n = 3 independent experiments. **C**, Localization of USP48 (green) in the nucleus of cultured GFAP (red)-positive rat glia. Scale bar, 20 µm. **D**, Single-cell RT-PCR detection of USP48 expression in a pyramidal neuron from a mouse hippocampus slice. PSD-95 was used as pyramidal neuron marker, GFAP was a glia marker, and GAPDH was used as a loading control. DNA ladder was loaded on the left lane. **E**, Co-localization of USP48 with the nuclear protein HDAC1 in HEK293T cells. Scale bar, 10 µm. **F, G**, Localization of Myc-USP48 in the nuclei of HEK293T (**F**) and HeLa cells (**G**). Nucleus was labeled with Hoechst (blue). Scale bars, 10 µm.

**Figure S2. Additional characterizations of nuclear localization of USP48 and mutants in HEK293T cells. A,** IF images of transfected HEK293T cells showing subcellular localization of Myc-tagged wild-type or mutant USP48 (green) relative to the nuclear Hoechst (blue) staining. **B**, Localization of GFP-fused USP48 and mutants (green) in HEK293T cells. **C**, IF images of transfected HEK293T cells showing cytoplasmic localization of Myc-USP48ΔNLS and GFP-USP48ΔNLS. Scale bars, 10 µm.

**Figure S3. A naturally exiting short isoform of USP48. A,** Sequence alignment of full-length mouse and human USP48 and USP48-S. hUSP48, full-length human USP48; mUSP48, full-length mouse USP48; hUSP48-S, a short isoform of human USP48; mUSP48-S, a short isoform of mouse USP48. NCBI ID number for each isoform is listed on the top. **B**, Schematic of the 484 amino acid mouse short isoform mUSP48-S that lacks 568 amino acids in the carboxyl terminus. **C**, Detection of the mUSP48-S PCR product from a mouse brain cortical cDNA library. **D**, IF images of transfected neurons showing that Myc-mUSP48-S and Flag-HA-hUSP48-S localized primarily in the cytoplasm and dendrites. Nuclei are stained with Hoechst. Scale bars, 20 μm.

**Figure S4. Characterizations of USP48 knockdown and knockout constructs. A, B**, IB analysis of knockdown efficiency in co-transfected HEK293T cells. n = 3 independent experiments. $^{**}p < 0.01$, $^{***}p < 0.001$, unpaired t tests. **C, D,** IF confirmation of USP48 knockdown efficiency in neurons transfected with shControl or shUSP48. Scale bar, 10 μm. n = 6 cells each group. $^{**}p < 0.01$, unpaired t test. **E**, IB confirmation of RNAi-resistant USP48 constructs (USP48r) in co-transfected HEK293T cells. **F**, IB analysis of CRISPR/Cas9-mediated USP48 knockout in stable HEK293T cells transfected with CRISPR or CRISPR USP48. Colonies of cells where full-length USP48 was completely deleted were identified by IB. **G**, IF images of USP48 WT and KO stable HEK293T cells confirming depletion of USP48 in KO cells. Scale bar, 10 μm. **H**, IF confirmation of USP48 knockout in neurons co-transfected with CRISPR or CRISPR USP48 and GFP. **I**, Quantification of **H**. n = 11 and 12 for CRISPR/GFP and CRISPR USP48/GFP, respectively. $^{***}p < 0.001$, unpaired t test.

**Figure S5. Additional data characterizing USP48 modulation of surface glutamate receptor levels and synaptic protein clustering. A**, Biotinylation assays of surface and total levels of GluA1, GluA2 and GluN2A in cultured neurons transfected with shControl or shUSP48. **B**, Quantification of relative protein levels from (**A**). n = 3 independent experiments. $^{**}p < 0.01$, unpaired t tests. **C**, Representative dendritic segments from neurons transfected with Myc-

GFP/GFP or Myc-USP48-S/GFP, showing immunostained Synapsin I clusters (red). Scale bar, 5 μm. **D**, Quantification of Synapsin I intensity from (**C**). n = 21 and 17 for Myc-GFP/GFP and Myc-USP48-S/GFP groups, respectively. **E**, Representative dendritic segments with immunostained PSD-95 clusters (blue) from neurons co-transfected with GFP, USP48/GFP, or C98A/GFP and DsRed-actin. Scale bar, 5 μm. **F**, Representative dendritic segments with PSD-95 clusters from neurons transfected with shControl or shUSP48. Scale bar, 5 μm. **G**, Quantifications of (**E**) and (**F**). n = 11, 11 and 15 dendritic branches for GFP, USP48/GFP and C98A/GFP, respectively. *P < 0.05, one-way ANOVA with post hoc Turkey's multiple comparison tests. n =10 and 8 dendritic branches for shControl and shUSP48, respectively. ***P < 0.001, unpaired t test.

**Figure S6. Additional experiments characterizing USP48 modulation of dendritic spines in vivo.** **A**, Representative low magnification image of a mouse hippocampal section (P32) infected with lentivirus expressing shUSP48/GFP injected at P18. Nuclei were labeled with Hoechst 33342. Scale bar, 350 μm. Doted box indicate where higher-magnification dendritic segments were sampled. **B**, Representative images of dendritic segments from CA1 neurons infected with GFP, USP48/GFP or C98A/GFP lentivirus. Scale bar, 20 μm. **C**, Representative images of dendrites from CA1 neurons infected with shControl/GFP or shUSP48/GFP lentivirus. Scale bar, 20 μm. **D**, Quantifications of total dendritic spine density from (**B)** and (**C**). n = 12 dendritic branches in each group. **P < 0.01, one-way ANOVA with post hoc Turkey's multiple comparison tests (left). ***P < 0.001, unpaired t test (right). **E**, Representative low magnification image of a mouse hippocampal section (P30) co-infected with AAVs expressing USP48/GFP injected at P0. Nuclei were labeled with Hoechst 33342. Scale bar, 350 μm. **F**, Representative dendritic branches at higher magnifications from CA3 neurons infected with GFP, USP48/GFP, or C98A/GFP AAVs. **G**, Representative dendritic branches at higher magnifications from neurons infected with shControl/GFP or shUSP48/GFP AAVs. Scale bars, 5 μm. **H**, Quantifications of total spine densities from (**F**) and (**G**). n = 12 in Control/GFP, USP48/GFP, and C98A/GFP group. **p < 0.01, one-way ANOVA with post hoc Turkey's multiple comparison tests. n = 9 and 10 in shControl and shUSP48 groups, respectively. ***p < 0.001, unpaired t test.

**Figure S7. Additional data supporting Figure 7. A**, Co-precipitation of Myc-USP48 and GFP-p65 in transfected HEK293T cells. IgG was used as a control. WCL, whole-cell lysates. **B**, Coomassie blue staining and IB assays of purified GST-GFP and GST-USP48 from HEK293T cells transfected with indicated plasmids. Arrowheads indicate purified GST-fused proteins. **C**, In vitro deubiquitination assay showing that purified GST-USP48 disassembled HA-Ub K48 polyUb chains. **D**, IB analysis of K48-linked polyubiquitination of p65 in Neuro2a cells transfected with shControl or shUSP48. **E**, IB analysis of p65 K48 polyubiquitination in USP48 knockout HEK293T stable cells. **F**, Luciferase reporter assay of NF-κB activation in neurons transfected with control, USP48, shControl or shUSP48 plasmids. n = 3 independent experiments. ***P < 0.001, unpaired t tests. **G**, Effects of USP48 knockdown on GFP-p65 levels in cultured neurons transfected with shControl or shUSP48 and GFP-p65. Scale bar, 20 μm. **H**, Quantification of GFP-p65 fluorescence level from (**G)**. n = 9 neurons for each group. ***P < 0.001, unpaired t test. **I, J,** IB analysis of p65 knockdown efficiency in cultured neurons. n = 3 independent experiments. ***p < 0.001, unpaired t test.
